# Synthetic gene circuits combining CRISPR interference and CRISPR activation in *E. coli*: importance of equal guide RNA binding affinities to avoid context-dependent effects

**DOI:** 10.1101/2023.06.13.544730

**Authors:** Içvara Barbier, Hadiastri Kusumawardhani, Lakshya Chauhan, Pradyumna Vinod Harlapur, Mohit Kumar Jolly, Yolanda Schaerli

**Author notes:** These authors contributed equally to the project (Hadiastri Kusumawardhani, Lakshya Chauhan).

## Abstract

Gene expression control based on CRISPR has emerged as a powerful approach for constructing synthetic gene circuits. While the use of CRISPR interference (CRISPRi) is already well-established in prokaryotic circuits, CRISPR activation (CRISPRa) is less mature and combination of the two in the same circuits is only just emerging. Here, we report that combining CRISPRi with SoxS-based CRISPRa in *Escherichia coli* can lead to context-dependent effects due to different affinities in the formation of CRISPRa and CRISPRi complexes, resulting in loss of predictable behaviour. We show that this effect can be avoided by using the same scaffold guide RNA structure for both complexes.

## Introduction

Synthetic biologists are building synthetic gene regulatory networks (GRNs) to decipher nature’s design principles (*1, 2*) and to provide new solutions to biomedical (*3*), agricultural (*4*), industrial (*5*), and environmental challenges (*6*). Despite impressive progress in constructing synthetic circuits (*7, 8*), the complexity of gene regulatory networks that has been achieved is still rather limited (*9–11*). Challenges to overcome include metabolic burden, resource competition, limited number of well-categorised parts, cross-talk between parts and context-dependent effects, leading to low modularity and scalability. While most synthetic circuits built so far have made use of protein transcription factors to regulate gene expression, clustered regularly interspaced short palindromic repeats (CRISPR)-based genetic regulation has the potential to address many of the current limitations (*12*). The advantages of CRISPR-based gene regulation tools compared to circuits based on protein transcription factors include decreased cross-talk between parts due to highly specific RNA-DNA interactions, reduced metabolic burden coming from protein production and straight-forward design of an unlimited number of orthogonal versions (*12*).

The strategies for transcriptional regulation based on CRISPR are known as CRISPR interference (CRISPRi) (*13*) and CRISPR activation (CRISPRa) (*14, 15*). In bacteria, the repression system uses a single guide RNA (sgRNA), which is composed of a target-specific sequence and a sequence that recruits a catalytically inactive version of Cas9 (dCas9). The complex is targeted to a promoter or a coding sequence to inhibit transcription. In contrast, for CRISPRa, dCas9 is targeted upstream of the promoter and it requires in addition an activator protein to recruit the RNA polymerase (*14, 15*). In prokaryotes, the activator protein can be directly fused to dCas9 (*16* –*21*) or alternatively, the sgRNA can be extended with a protein-recruiting RNA scaffold that recruits the transcriptional activator (*22–26*). Probably the best characterized bacterial CRISPRa system is based on a so-called scaffold RNA (scRNA) where the sgRNA is modified to include a 3’ MS2 hairpin. This hairpin recruits the MS2 coat protein (MCP) that is fused to the transcriptional activator SoxS (*22* –*25*).

We recently showed that CRISPRi can be used to build dynamic and multistable synthetic circuits (*28*). Extending such circuits with CRISPRa has the potential to further increase the complexity of synthetic gene regulation programs. Here, we show that combination of CRISPRi with CRISPRa based on scRNA and SoxS can lead to strong context-dependent effects. Specifically, the strength of activation mediated by scRNA was strongly influenced by the concurrent expression of a sgRNA. We hypothesized that this phenomenon was caused by sgRNA and scRNA competing for the limited pool of dCas9 and their differential affinities to dCas9. This hypothesis was supported by a mathematical model. The model also suggested ways to circumvent this problem. We then experimentally reduced this context-dependent effect by using scRNAs for both repression and activation and thus improved the predictability of synthetic CRISPRa/i circuits.

## Results

### Implementing CRISPRa

We implemented CRISPRa using our previously developed plasmid architecture and cloning strategy (*29*), which we had employed to construct multistable and dynamic CRISPRi-based circuits (*28*). Our CRISPRi system is composed of two plasmids: The first plasmid (colA ori) harbors the designed circuit with up to three nodes of which one is arabinose-inducible via a pBAD promoter. From the second plasmid (CDF ori) we constitutively express dCas9 and Csy4. We use Csy4 RNase-processing to release parts that are transcribed together in the same operon to act independently once transcribed, such as sgRNAs/scRNAs for CRISPRi/a and mRNAs coding for a fluorescent reporter.

For CRISPRa, we added a third plasmid (pBR322 ori) constitutively expressing (promoter J23119) MCP-SoxS (carrying mutations R93A+S101A) (*22*). We started with a two nodes circuit (Figure 1A) with the first node containing an arabinose-inducible scRNA (version b2 (*23*)) guiding dCas9 and MCP-SoxS to bind and subsequently activate the second node containing a weak promoter (J23117) upstream of a green fluorescent protein (GFP) reporter. As CRISPRa is very sensitive to the distance between the target site and the transcriptional start site (TSS), we used a sequence ranging from the target site to the TSS previously shown to work (J306 and J3 region) (*22*). We confirmed that the 81 bp distance from TSS is ideal for activation (Figure S1) and we obtained around 20-fold activation compared to the off-target control (Figures S1 and S2).

**Figure 1:**
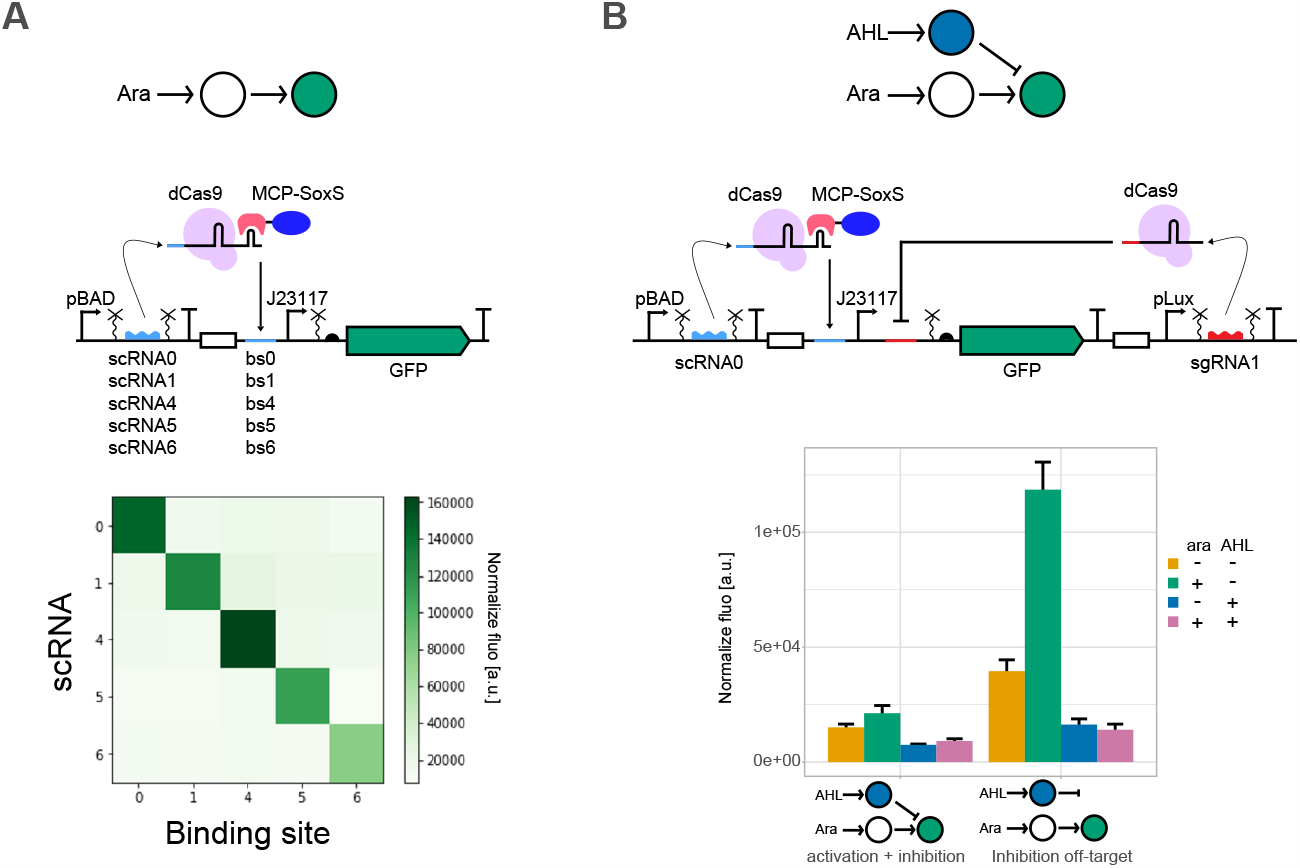
CRISPRa orthogonality and combination with CRISPRi. **A**. CRISPRa orthogonality. Top: Schematic representation of the circuit and details of the circuit design. Symbols according SBOL standard (27). Bottom: orthogonality heatmap of GFP fluorescence normalized by the absorbance at 0.2% arabinose with different combinations of scRNAs and binding sites as represented in the figure. **B**. Combination of CRISPRa and CRISPRi. Top: Details of the circuit design. Middle: Schematic representation of the circuit. Bottom: bar plots represent the GFP fluorescence normalized by the absorbance in the absence or presence of arabinose (0 and 0.2%) and AHL (0 and 0.1μM). Binding site number 2 was used for off-target inhibition. Mean and s.d. represent three biological replicates.

Once we had successfully integrated CRISPRa into our framework, we created an orthogonal library of scRNAs (Figure 1A). We added the target-specific sequences of 6 previously characterised orthogonal sgRNAs (*28, 30*) to the scRNA scaffold and replaced the sequence 81 nt upstream of the TSS with the corresponding binding sites. Four (numbers 1, 4, 5 and 6) out of the six tested constructs resulted in detectable activation (Figure S2). Together with the original scRNA (number 0) we tested their orthogonality (Figure 1A). This analysis confirmed that we observe activation only when the scRNA and binding site pairs match. We noticed that the non-induced controls of matching pairs resulted in higher green fluorescence levels than non-matching pairs of scRNA and binding site (Figure S2). We attribute this to the previously reported leakiness of our pBAD promoter (*28*).

### Combining CRISPRa and CRISPRi

Next, we combined CRISPRa and CRISPRi in the same circuit. We added a third node to our activation circuit, which in presence of AHL produces a sgRNA complementary to a binding site placed downstream of the promoter repressing the expression of GFP in the second node (Figure 1B). We expected that GFP expression increases in presence of arabinose and decreases in presence of AHL. In the presence of both inducers, the expression depends on the relative strength of the the two opposing inputs, but as the binding site for the CRISPRi complex is downstream of the promoter, we expected the repression to be dominant. However, the circuit showed a very low level of activation in presence of arabinose only. In our off-target control (orthogonal binding site 2 instead of binding site 1) for inhibition, we observed the expected activation with scRNAs induction, which might indicate that leaky expression of the sgRNA is enough to repress the activation in the full circuit. Moreover, in the off-target control we noticed a strong repression upon induction of the sgRNA, even though the sgRNA should not repress. These results let us to hypothesize that the sgRNA competes with the scRNA for dCas9, with an advantage for the CRISPRi complex.

### Model suggests differential affinities of scRNA and sgRNA are a problem

To test our hypothesis, we adapted a qualitative mathematical model (*31*) describing the transcription of scRNA and sgRNAs, the formation of the CRISPRa and CRISPRi complexes, their binding to DNA and subsequent transcriptional activation or repression, respectively (Figure 2A, see methods). Then, we varied the key parameters of CRISPRa/i complex formation (*k*_*i*_, *k*_*j*_) and binding of the complexes to DNA (*q*_*i*_,*q*_*j*_) (Figure 2B-E). We found that we could reproduce our experimental finding of Figure 1 when CRISPRi parameters *k*_*j*_ and *q*_*j*_ are of several orders of magnitude bigger than their CRISPRa counterparts (*k*_*i*_ and *q*_*i*_) and when we have some leaky expression of sg/scRNAs (Figure 2E). This suggests that the scRNA binds weaker to dCas9 than the sgRNA and the CRISPRa complex binds weaker to DNA than the CRISPRi complex. Thus, the model supported the hypothesis that scRNA and sgRNA compete for the pool of available dCas9.

**Figure 2:**
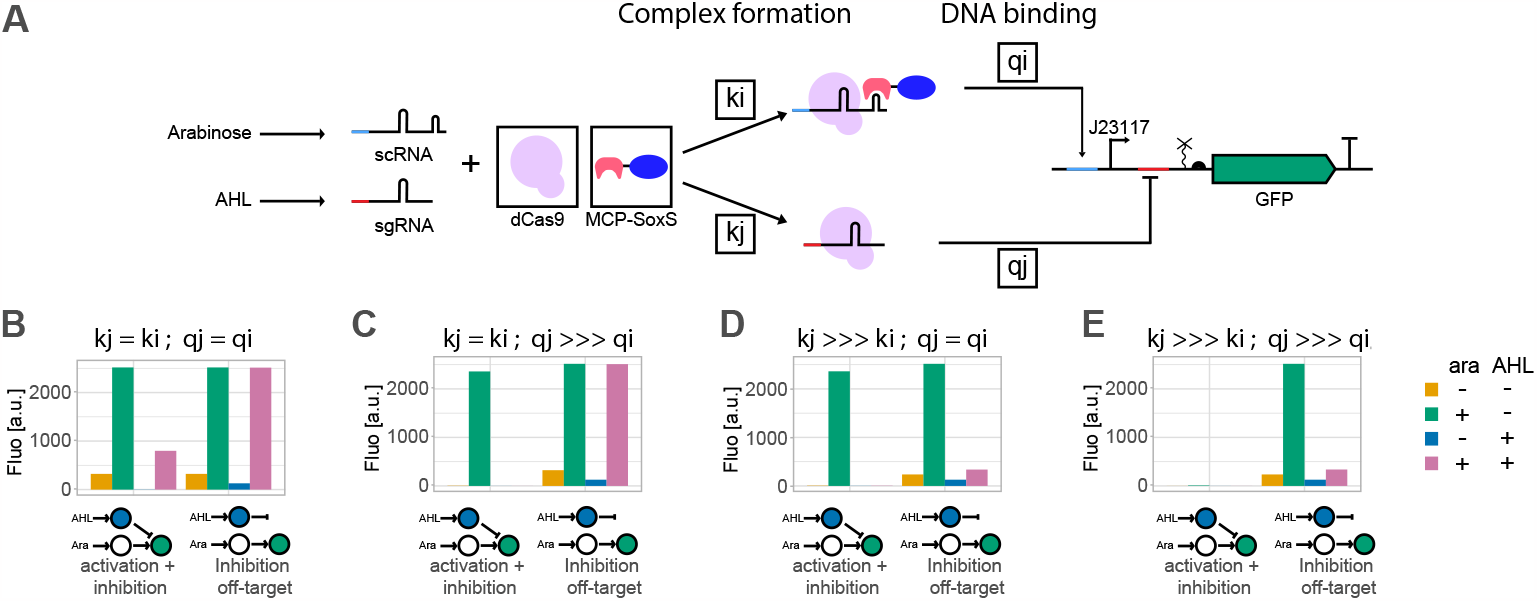
Model suggests that differential affinities of scRNA and sgRNA are a problem. **A**. Schematic representation of the Quasi-Steady State Approximations model. The different elements changed in the analysis are indicated with black boxes: quantity of dCas9 and MCP-SoxS, values of complex formation parameters (ki, kj) and values of complex binding to DNA (qi, qj). See methods for detailed equations. **B-E**. Qualitative model of GFP intensity with or without arabinose and AHL induction (0,10) and different values of kj and qj (1 and 100000), ki and qi are fixed at 1. For the off-target controls, qj is set to 0. Situation E resembles most our experimental data. Situations B and C show the desirable behaviour.

Thereby, the model put forward a potential solution to achieve the expected behaviour: to ensure the complex formation rates are similar for CRISPRa and CRISPRi (*k*_*i,j*_) (Figure 2B,2C). The DNA binding affinities (*q*_*i,j*_) are less important because dCas9-sgRNA and dCas9-scRNA-MCP-SoxS complexes do not bind to the same binding sites. However, when the complex formation rates are unequal (*k*_*j*_ *>>> k*_*i*_), a very strong binding of the inhibiting CRISPRi complex *q*_*j*_ compared to the activating CRISPRa complex *q*_*i*_ leads to an absence of activation in the on-target circuit (Figure 2E), while similar DNA binding rates *q*_*j*_ = *q*_*i*_ allow for proper activation in the on-target circuit but still show an incorrect off-target behaviour (Figure 2D).

We also investigated the influence of dCas9 and MCP-SoxS quantities in our model (Figure S3). Increasing the amount of MCP-SoxS helps to increase the activation level but comes at a price of increased leaky activation in the absence of the inducer (arabinose). Increasing the amount of dCas9 allows the correct behavior of the off-target control but not of the on-target circuit. We hypothesise that the dCas9 increase does not rescue the behavior because of the genetic configuration, paired with the use of slightly leaky promoters: the binding site for inhibition is downstream of the activation binding site. Thus, if both CRISPRa and CRISPRi complexes are bound, transcription is repressed and the leaky expression of sgRNA and a high concentration of dCas9 is sufficient to do so even in the absence of AHL inducer. Therefore, increasing the total amount of dCas9 is not predicted to recover the correct behavior of our circuits. Anyway, high expression levels of dCas9 are known to be toxic to *E. coli* cells (*32*) and MCP-SoxS expression was already maximized with a strong promoter on a high-copy plasmid. Therefore, equalizing the complex formation rates promised to be the most promising approach.

### Using scRNA for CRISPRi and CRISPRa restores the function

We thus set out to test the model predictions and attempted to make the complex formation rates similar for CRISPRa and CRISPRi. We first tested if truncating the sgRNA by 4 bp (sgRNAt4) would lead to the desired behavior (Figure S4A). Truncated sgRNAs-dCas9 complexes display weaker repression than their full-length counter-parts (*13*), but the DNA binding of the CRISPR complex is similar as with a full-length sgRNA (*33, 34*). We observed the correct behavior for the off-target control but still almost no activation when combined with the on-site repression (Figure S4A). This behavior can be reproduced in our model when sgRNAt4 binds weaker to dCas9 than sgRNA but still stronger than scRNA while DNA binding affinities of sgRNAt4 and sgRNA4 are similar (Figure S4B).

Next, we used scRNA instead of sgRNA also for the inhibition complex. As observed for other CRISPRa systems (*19, 20*), if we directed dCas9-scRNA-MCP-SoxS downstream of a promoter we observed inhibition, rather than activation (Figure S5). We thus rebuilt the circuits in Figure 1, but this time with scRNAs for both CRISPRa and CRISPRi (Figure 2). Now we observed a good level of activation in the presence of arabinose in our on-target circuit and no inhibition with AHL induction in the off-target control. This data demonstrates that in agreement with our model, ensuring similar complex formation rates allows for the correct functioning of combined CRISPRa and CRISPRi circuits. Therefore, using the same scRNAs for both inhibition and activation is a straightforward way to obtain the expected circuit function.

### Cascade circuits

Encouraged by the predictable behavior when using scRNA for both CRISPRa and CRISPRi complexes, we proceeded to combine CRISPRa and CRISPRi in a cascade circuit. Here, the first node is induced by arabinose and represses the second node (containing a mKate reporter) that activates the third node encoding a GFP reporter (Figure 3A). This circuit also behaved as expected: we observed expression of GFP and mKate in absence of arabinose and their level decreased upon addition of arabinose. In addition, we built three off-target controls. All measurements of the controls agreed with our expectations, while using sgRNA for inhibition led again to an incorrect behaviour of the off-target controls (Figure S6). We thus demonstrated, that CRISPRa and CRISPRi can be successfully combined, but special attention has to be paid to different affinities in RNA-dCas9 complex formation and DNA binding.

**Figure 3:**
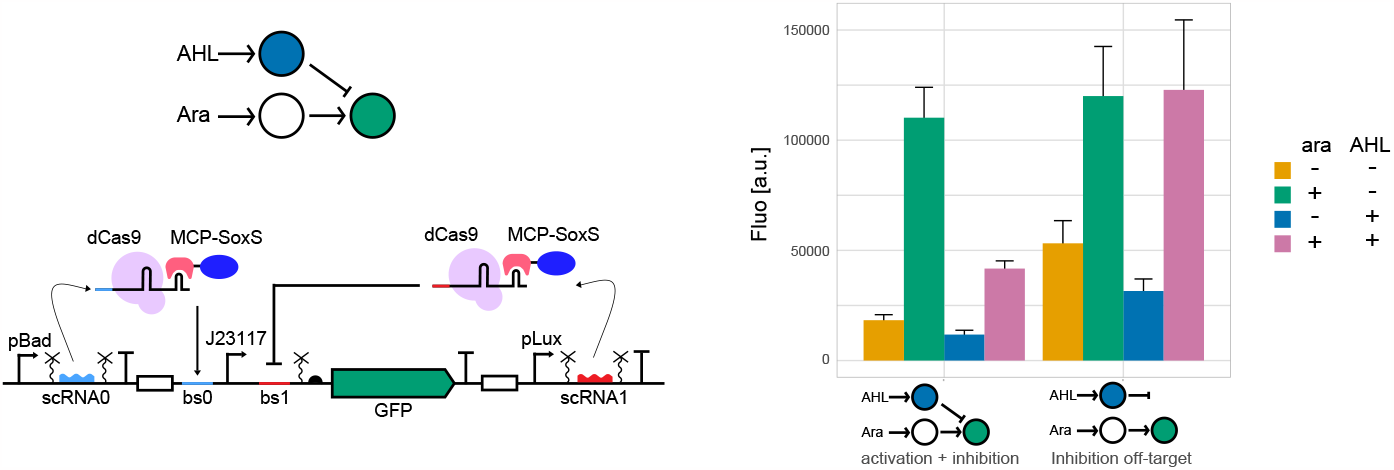
Using scaffold RNA in both CRISPRa and CRISPRi restores predictable circuit behaviour. Left: Details of the circuit design and schematic representation of the circuit. Right: bar plots represent the GFP fluorescence in absence or presence of arabinose (0 and 0.2%) and AHL (0 and 0.1μM). Binding site number 2 was used for off-target inhibition. Mean and s.d. represent three biological replicates.

**Figure 4:**
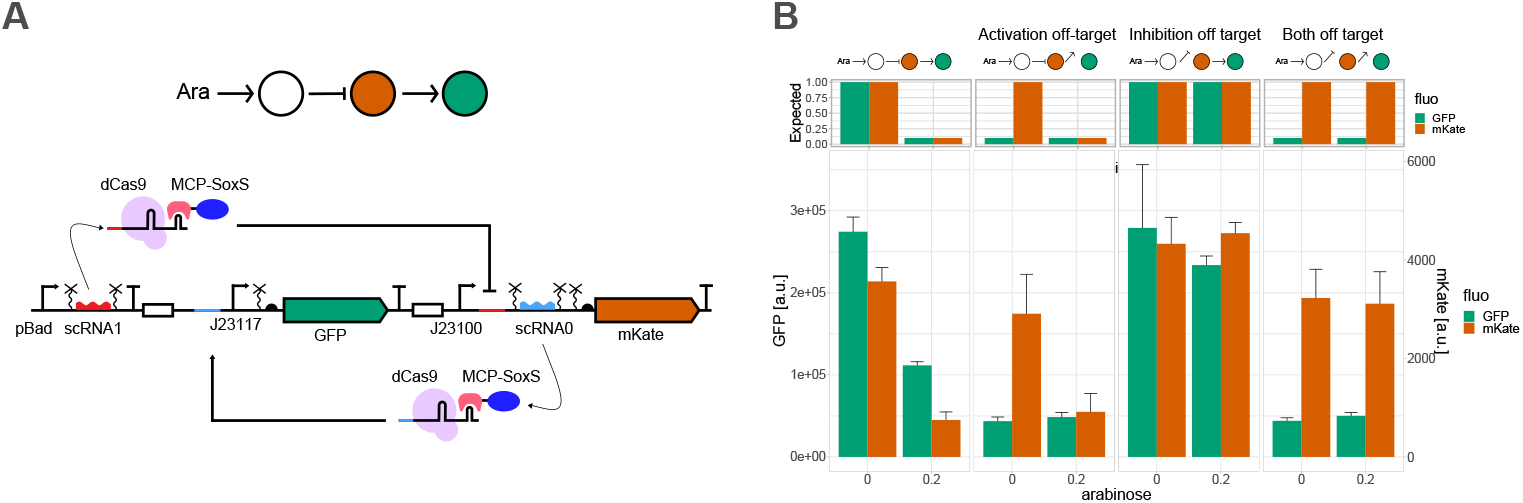
Combination of CRISPRi and CRISPRa in a cascade circuit. **A**. Details of the circuit design and schematic representation of the circuit. **B**. Bar plot representing the GFP and mKate fluorescences of different circuits with different off-target controls. The expected behaviours are illustrated at the top. Mean and s.d. represent three biological replicates.

## Discussion

In this work, we successfully implemented the SoxS-based CRISPRa system first described by Fontana *et al*. (*22*) into our framework (*28, 35*). When we combined CRISPRa with CRISPRi, we observed a strong affinity competition for dCas9 leading to weak activation and incorrect circuit function of the off-target controls. This can lead to an undesirable coupling among circuit branches that theoretically should act orthogonal. Guided by mathematical modelling, we managed to avoid this problem and obtained the circuits’ correct function by using the same RNA (i.e. scRNA) for CRISPRa and CRISPRi resulting in the same complex affinities for activation and inhibition.

Competition for transcriptional and translational resources is a well known issue in engineering synthetic circuits (*9, 36*). Moreover, it has also been shown that expressing simultaneously multiple sgRNAs that compete for the same limited pool of dCas9 can lead to unwanted outcomes (*31, 37*). Here, we describe yet a different problem related to dCas9 resource competition when combining CRISPRi (using sgRNA) and CRISPRa (using scRNA) in one bacterial cell: The different guide RNAs have different affinities for dCas9 and thus competition hampers correct network function. Previous work by Tickman and colleagues combined CRISPRi and CRISPRa in different circuits such as cascades and incoherent feed-forward circuits in cell-free extract and in *E. coli* (*24*). However, they did not report a resource competition between the two systems. It might be that they had conditions with very tight sgRNA production and high dCas9 concentration where competition was minimized.

While building synthetic circuits with CRISPRi and CRISPRa is rather new in prokaryotes, the combination of CRISPRa and CRISPRi had been used to control host genes in yeast and mammalian cells (*38* –*43*). In these systems, the competition between CRISPRa and CRISPRi paths was not observed, as they either used orthogonal Cas proteins (*38* –*40*) or scRNAs for CRISPRa and CRISPRi as both functions require regulator domains (*41* –*43*).

Here, we present a simple solution for the encountered problem in *E. coli* by also using scRNAs for both CRISPRi and CRISPRa. Our model suggests that simply increasing the dCas9 pool is not able to restore the correct function in our circuits (Figure S3). This is due to the leaky production of sgRNA and the dominant effect of repression over activation caused by the circuit architecture. While increasing the concentration of dCas9 may help in other cases of dCas9 competition, overproduction of dCas9 can lead to toxicity, reduced growth rates and morphological defects (*32, 37*). Reported solutions to address this problem include the use of a non-toxic variant of dCas9 (*37*) and regulated production of dCas9 adapting to the current circuit load (*44*). For future work it would be interesting to test these approaches with circuits combining CRISPRi and CRISPRa. We hope our work paves the way for building more complex bacterial CRISPRa/i circuits and their applications for studying the function of native genes, cellular reprogramming, and metabolic engineering (*45* –*48*)

## Materials and methods

### Plasmids construction

Circuits were built as previously described (*29*). The different parts contain prefix (CAGCCTGCGGTCCGG) and suffix (TCGCTGGGACGCCCG) sequences (*49*), which can be PCR amplified (Phanta Max Super-Fidelity DNA Polymerase, Vazyme) with a set of primers (ordered from Microsynth or Sigma-Aldrich) to add a unique variable linker. Backbones were linearized with PCR or with restriction enzymes (NEB, 1h at 37°C). PCR-amplified or digested products were purified (Monarch PCR & DNA Cleanup Kit, NEB). Then, the parts were assembled with Gibson assembly (NEBuilder HiFi DNA Assembly Master Mix from NEB, 1h at 50°C) with the linkers providing sequence overlaps. Finally, 1 *μl* of the assembly mixes were transformed into competent cells (NEB5*α* cell) by electroporation and plated on LB agar plate containing appropriate antibiotics (50*μg/l* kanamycin, 100*μg/l* ampicillin or 50*μg/L* spectomycin). The obtained plasmids were sequenced (Microsynth) to confirm that they contained the correct constructs. Complete plasmid sequences are provided as supporting information.

### Fluorescence measurements

Plasmids were co-transformed into Mk01 *E. coli* cells (*50*). Single colonies were incubated in 200*μl* of EZ medium with 0.4% glycerol as carbon source (Teknova) with appropriate antibiotics (25*μg/L* kanamycin, 50*μg/L* ampicilin and 25*μg/l* spectomycin) at 37°C, 200 RPM for 4 to 5 hours. Then, cells were diluted to 0.05 OD600 in 96-well CytoOne plate (Starlab) with or without inducers, as indicated in the figures. Plates were incubated at 37 °C with double-orbital shaking (Synergy H1 microplate reader, Biotek, running Gen5 3.04 software). Fluorescence was determined after 16 h with 479 nm excitation and 520 nm emission for GFP and 588 nm excitation and 633 nm emission for mKate2. Fluorescence values were normalized by division with absorbance at 600nm. Subsequent data were analyzed and visualized with R.

### Modelling

The model is based on mass action law kinetics and Quasi-Steady State Approximations (QSSA) accounting for various molecular steps and constraints such as copy number of plasmids and steady-state protein levels. Moreover, we assume that the binding of sgRNA/scRNA in inhibiting and activating edges are independent of each other. The model is explained below, where the set of equations with subscript *i* refer to activation due to arabinose, and *j* refers to inhibition due to AHL. The parameters used are explained in Table 1. The code is available at https://github.com/SchaerliLab/CRISPRa-i-circuits

**Table 1:** Parameters’ description and values.

| Parameter | Description | Value | Unit |
| --- | --- | --- | --- |
| $\delta_{RNA}$ | Degradation constant for first order degradation of guide RNA | 10.8 | hr-1 |
| $\gamma_{i,j}$ | Transcription rates of guide RNA/scRNA (competing guide RNAs) | 530.1 | nMhr-1 |
| $K_j$ | Transcription rate due to leaky promoter | 5 | hr-1 |
| $K_i$ | Transcription rate from dCas9 bound promoter (activation) | 100 | hr-1 |
| $\delta_{GFP}$ | Degradation constant for first order degradation of GFP RNA | 1.176 | hr-1 |
| $D_{i,j}$ | Concentration of free target DNA binding sites for scRNA (i) or sgRNA (j) | variable | nM |
| $D_{i,j,total}$ | Total concentration of target DNA binding sites for scRNA (i) or sgRNA (j) | 30 | nM |
| $d_{total}$ | Total quantity of dCas9 | 100 or 10000 (figure 2F) | |
| $m_{total}$ | Total quantity of MCP-SoxS | 100 | |
| $k_i$ | scRNA ($x_i$), dCas9 and MCP-SoxS complex formation affinity | 1 | |
| $k_j$ | sgRNA ($x_j$) and dCas9 complex formation affinity | variable (1 - 100000) | |
| $q_i$ | DNA binding affinity of the CRISPRa complex | 1 | |
| $q_j$ | DNA binding affinity of CRISPRi complex | 1 or 100000 | |
| $x_i$ | concentration of scRNA | variable | |
| $x_j$ | concentration of sgRNA | variable | |

Rate of change of scRNA and sgRNA from arabinose and AHL induction respectively:

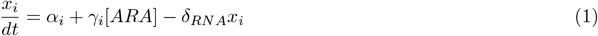

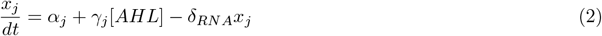

Formation of CRISPRa and CRISPRi complexes (lacking DNA binding sites):

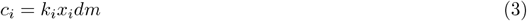

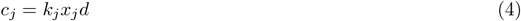

Formation of CRISPRa and CRISPRi complex with DNA binding sites:

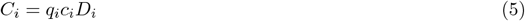

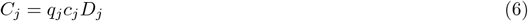

Constraints on DNA binding sites,

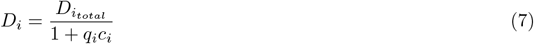

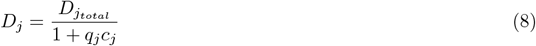

Net total available sites are the plasmid copy number and are a sum of free DNA binding sites and sites in CRISPRa/i complexes.

Rate of production of GFP from activation due to CRISPRa, leaky expression and first order RNA degradation

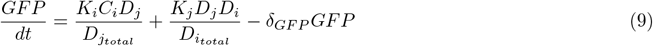

In our experimental system, we would expect the maximum production of GFP when CRISPRa is causing proper activation of the promoter (i.e. complex *C*_*i*_) and there is no inhibition due to CRISPRi (complex *D*_*j*_). In our model, we implement this using principles of conditional probability and obtain the net number of such sites as 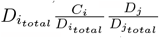 which multiplied by the production rate of GFP RNA gives us the first term of equation (9). We also expect leaky production of GFP RNA when both CRISPRa and CRISPRi are inactive. The total number of such sites available are 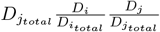 which is multiplied by the rate of basal expression (*K*_i_) to capture leaky expression in our model. We assume that any presence of CRISPRi complex will lead to a complete block of transcription. It is important to note that the values of 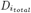 and 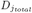 are the same in our system as both our sites are present on the same plasmid.

Constraint equations for dCas9 and SoxS proteins

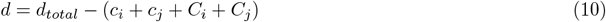

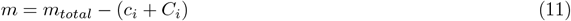

In the equations above, *x*_*i,j*_ are the concentration of scRNA(i) and sgRNA(j), *c*_*i*_ the scRNA, dCas9 and MCP-SoxS complex, *c*_*j*_ the sgRNA and dCas9 complex, *C*_*i,j*_ the activation or inhibition complex bound to DNA, *D*_*i,j*_ the amount of free DNA binding sites, *d* the concentration of free dCas9 and *m* the concentration of free MCP-SoxS. The descriptions and values of the different parameters are summarized in Table 1. Due to the lack of studies on the biochemical properties of CRISPRi and CRISPRa, arbitrary values were chosen for all parameters not available in the literature. The model was updated with a simplistic Euclidean update and brent equation solver in python3.

## Supporting information

Plasmid sequences

## Author contributions

I.B, H.K and L.C performed the experiments and analyzed the research, I.B and Y.S wrote the manuscript, L.C. and P.V.H. performed the modelling, M.K.J and Y.S. supervised the project. Y.S. acquired funding.

## Acknowledgments

This work was funded by the Swiss National Science Foundation (grants 31003A 175608 and 310030 200532 awarded to Y.S). We thank Louise Martin and Lucie Gilli é ron for their help with cloning.

