## Supporting Figures for "Synthetic gene circuits combining CRISPR interference and CRISPR activation in *E. coli*: importance of equal guide RNA binding affinities to avoid context-dependent effects"

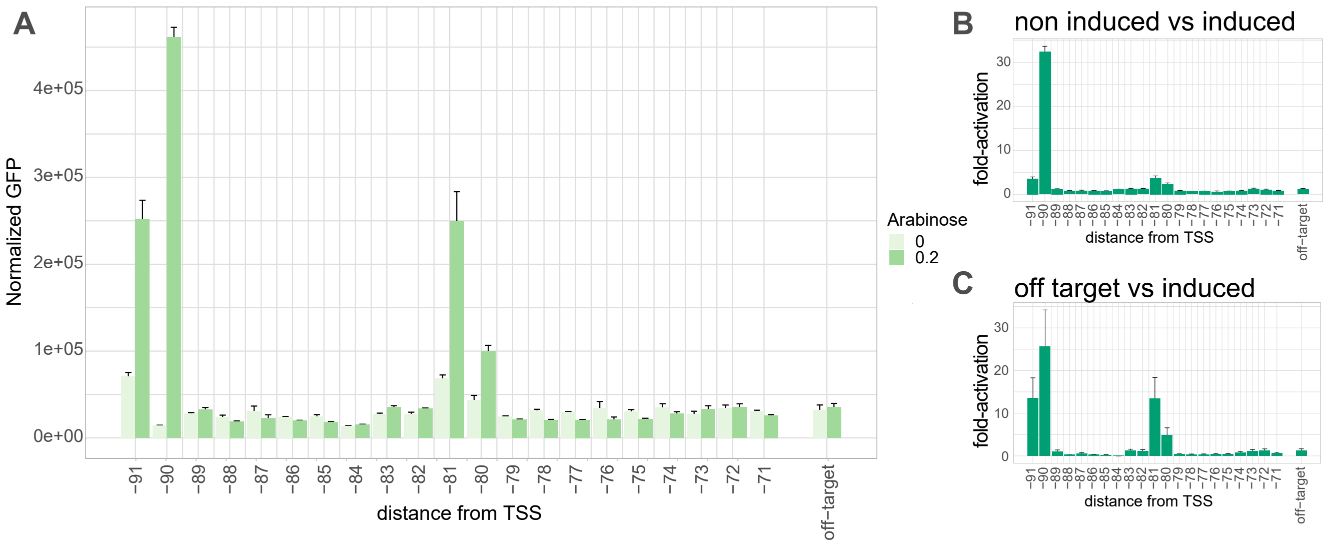


**Figure S1: Activation is sensitive to the distance to the transcriptional start site (TSS)**
***A.*** *Normalized green fluorescence at indicated arabinose concentrations. Different binding site position upstream of the TSS were tested as indicated in the x axis.* ***B.*** *Fold-activation at 0.2% arabinose compared to fluorescence level at 0% arabinose.* ***C.*** *Fold-activation at 0.2% arabinose compared to off-target construct (scRNA 1 - binding site 4) at 0% arabinose. Mean and s.d. represent three biological replicates.*

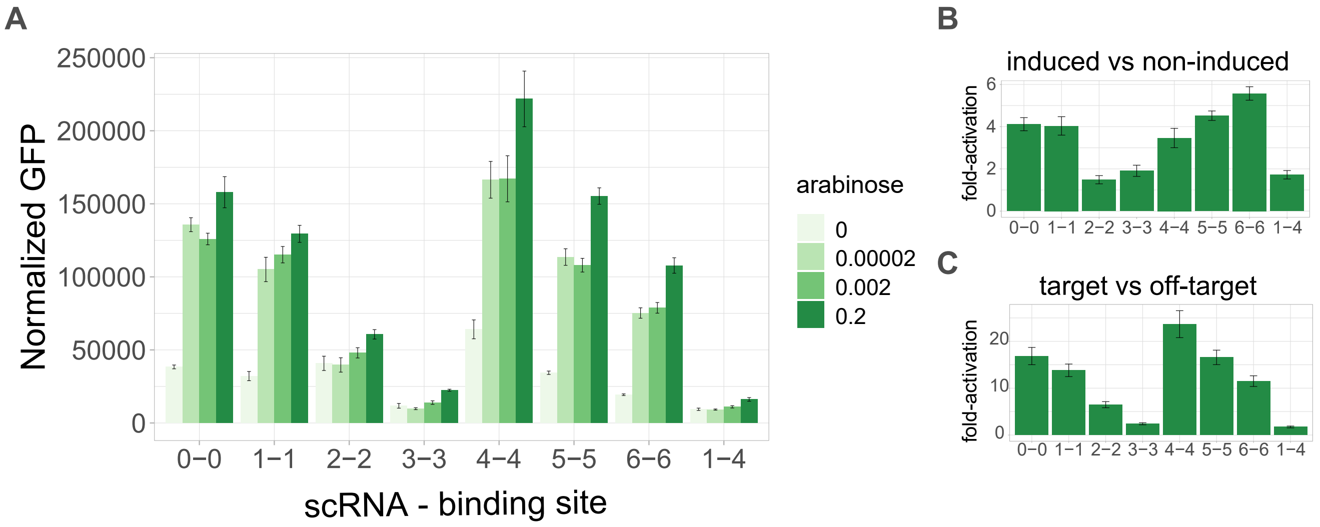


**Figure S2: Induction of CRISPRa at different arabinose concentrations**
***A.*** *Normalized green fluorescence at indicated arabinose concentrations. Different combinations of scRNAs and binding sites were tested as indicated in the x axis.* ***B.*** *Fold-activation at 0.2% arabinose compared to fluorescence level at 0% arabinose.* ***C.*** *Fold-activation at 0.2% arabinose compared to off-target construct (scRNA 1 - binding site 4) at 0% arabinose. Mean and s.d. of three biological replicates.*

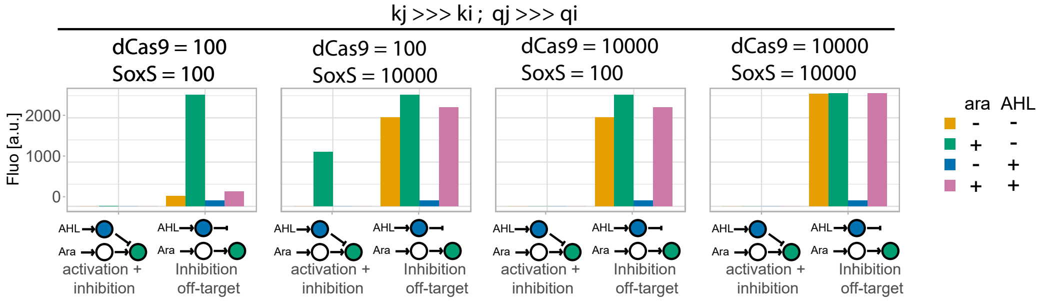


**Figure S3: Increasing dCas9 and SoxS quantities do not lead to correct function.**
*Qualitative model of GFP intensity with or without arabinose and AHL induction (0,10) and different values of MCP-SoxS and dCas9. kj and qj are fixed at 100000, ki and qi are fixed at 1. For the off-target controls, qj is set to 0.*

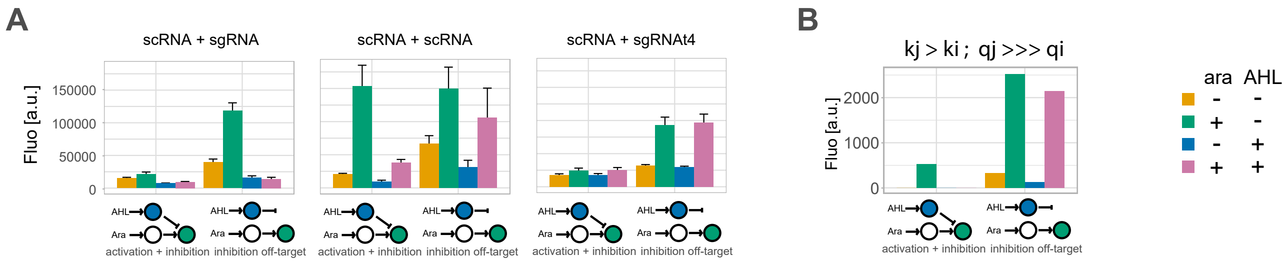


**Figure S4: CRISPRa/i induction with different single guide RNAs**
***A.*** *Bar plots at 0 or maximum concentration of AHL (0.1 $\mu M$) and arabinose (0.2%). Data in A and B are the same. Mean and s.d. represent three biological replicates.* ***B.*** *Modeling as in figure 2 but with $qj = 100000$ and $kj = 1000$*

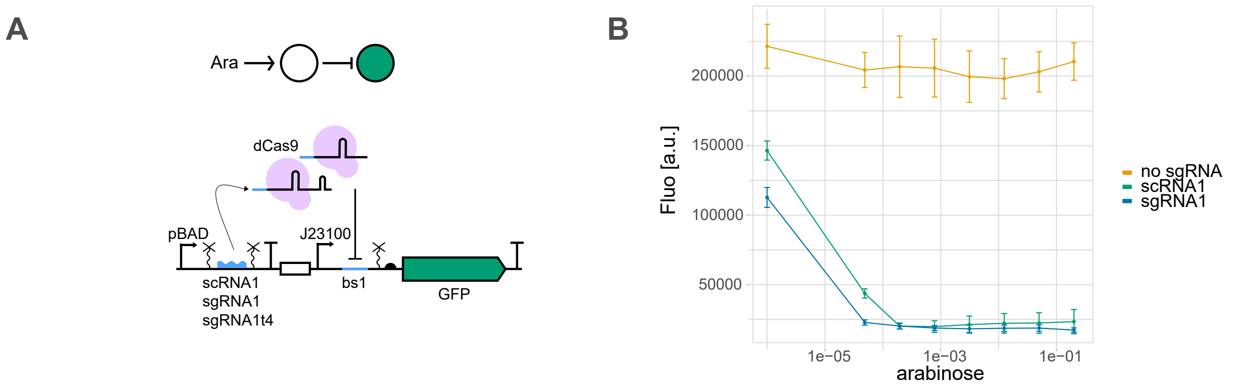


**Figure S5: Inhibition with sgRNA and scRNA**
***A.*** *Details of the circuit design and schematic representation of the circuit.* ***B.*** *GFP fluorescence in presence of different concentrations of arabinose. Mean and s.d. represent three biological replicates.*

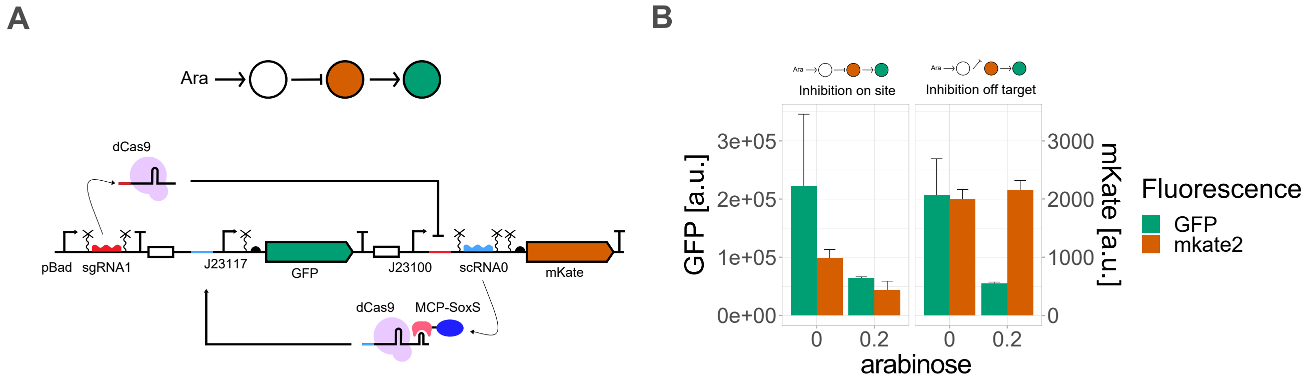


**Figure S6: Cascade circuit with sgRNA and scRNA leads to unpredicted behavior of the off-target control**
***A.*** *Details of the circuit design and schematic representation of the circuit. It is the same as in Figure 4, but instead of using scRNA1, we used here sgRNA1* ***B.*** *Bar plots represent the GFP and mKate fluorescences of the circuit and of an off-target control in absence (0%) or presence of arabinose (0.2%). Mean and s.d. represent three biological replicates.*
